# BIOSPONGES EMBEDDED WITH GDNF ENHANCE NEUROMUSCULAR RECOVERY FOLLOWING VOLUMETRIC MUSCLE LOSS

**DOI:** 10.64898/2025.12.10.693478

**Authors:** Jamshid Tadiwala, Connor Tobo, Kevin D. Sekerak, Rebecca Sheetz, Amelia Ridolfo, Madhushika Elabada Gamage, Elif G Ertugral, Paul Jelliss, Matthew D. Wood, Chandrasekhar R Kothapalli, Koyal Garg

**Affiliations:** Department of Biomedical Engineering, School of Science and Engineering, Saint Louis University, Saint Louis, MO; Department of Chemistry, Saint Louis University, Saint Louis, MO; Department of Surgery, Washington University in St. Louis School of Medicine, Saint Louis, MO; Department of Chemical and Biomedical Engineering, Cleveland State University, Cleveland, OH

## Abstract

Skeletal muscle cannot regenerate after volumetric muscle loss (VML), a traumatic injury defined as the loss of > 20% of a muscle’s mass. VML directly reduces the number of myofibers and causes axonal degeneration of nerves, resulting in reduced muscle function and impaired neuromuscular junctions (NMJs). Biosponge (BSG) scaffolds, composed of gelatin, collagen, and laminin-111, have been shown to improve muscle mass, cross-sectional area, and myofiber number following VML. However, improvements in NMJ quantity were not observed. Glial cell line-derived neurotrophic factor (GDNF) is a growth factor that enhances motor unit survival and neurite outgrowth. In this work, BSG scaffolds were electrostatically coupled with GDNF via gelatin nanoparticles (GNPs) to support myofiber regeneration and preserve NMJs post-VML in a rodent model. *In vitro* determination of release kinetics revealed an initial burst release of surface bound GDNF with almost an equivalent amount of electrostatically bound GDNF retained within the BSG post 1 week of incubation at 37°C in phosphate buffered saline (PBS). To create the VML injury in male Lewis rats (10-12 weeks old), ∼20% of the muscle mass was removed from the tibialis anterior (TA) muscle of both hindlimbs. Relative to BSG+GNP alone, treatment with BSG+GNP+GDNF showed a significant increase (∼25%) in peak isometric torque at 6 weeks post-injury. Qualitative and quantitative histological analysis of NMJs revealed an enhanced overlap between pre- and post-synaptic structures in the BSG+GNP+GDNF group. Additionally, the incorporation of GDNF slowed BSG remodeling and degradation. Overall, these results suggest that the BSG-mediated delivery of GDNF is an effective strategy for mitigating NMJ loss and enhancing muscle recovery following VML.

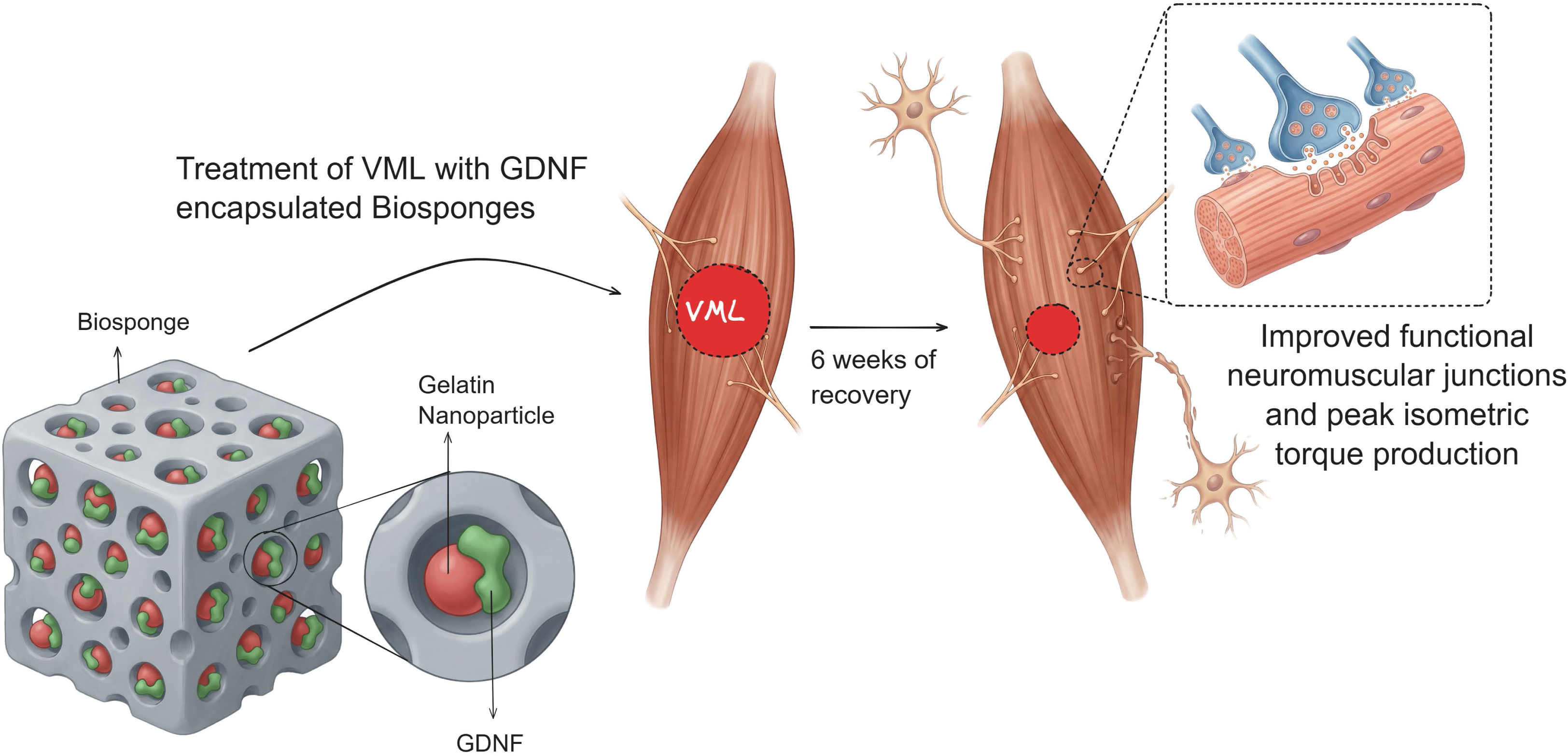

Graphical Abstract Tadiwala et al., 2025 Biosponges embedded with GDNF promote neuromuscular recovery following volumetric muscle loss.

## 1. Introduction

Skeletal muscle tissue cannot regenerate after volumetric muscle loss (VML), a traumatic injury defined as the loss of >20% of a muscle’s mass. Preclinical animal models of VML injury indicate that a relatively small VML injury (10-20% loss of muscle) can disproportionately reduce function (30 -90% peak isometric torque deficit) [1, 2]. VML directly reduces the number of myofibers and causes axotomy of nerves, resulting in reduced muscle function and fewer functional neuromuscular junctions (NMJs). In a recent study, a ∼20% VML defect resulted in chronic axotomy of ∼69% of the motor neurons innervating that muscle [3]. The reinnervation of tissue-engineered skeletal muscle (or denervated muscle) is a slow and extensive process that may require >3 months for maximal recovery [4, 5]. Together, these observations may provide the basis for the disproportionately higher loss of muscle function.

Recent work has also shown a continuous increase in denervation of VML-injured muscles [6]. Within 3 days of injury, denervation increased by 10%, but further escalated to 22% and 32% on days 21 and 48 post-injury, respectively. These changes were accompanied by a progressive increase in irregular morphological characteristics for NMJs (e.g., polyinnervation, axon sprouting, and endplate fragmentation). A similar progression of worsening pathology has been reported in both aged [7] and dystrophic [8] skeletal muscles.

Our lab has developed biosponge (BSG) scaffolds composed of gelatin, collagen, and laminin-111, which have improved muscle mass, function, cross-sectional area, and myofiber number following VML [9–12]. However, improvements in NMJ quantity were not observed. Glial cell line-derived neurotrophic factor (GDNF) is a growth factor that enhances motor unit survival and neurite outgrowth [13–18]. This study aimed to evaluate how incorporating GDNF into the structure of a BSG scaffold would improve the quantity and morphology of NMJs, as well as muscle function post-VML. We postulated that electrostatic conjugation of GDNF to a biomaterial carrier would protect and stabilize the growth factor while enabling sustained release through controlled dissociation. GDNF plays a crucial role in maintaining NMJ health through multiple mechanisms [14], including regulating motor neuron survival, protecting motor neurons from chronic degeneration, and rescuing them from axotomy-induced cell death. While GDNF expression is induced and upregulated in denervation models up to 48 hours post-injury, this endogenous response is not sustained enough for long-term or complete recovery [19].

We hypothesized that BSG scaffolds encapsulating GDNF-decorated gelatin nanoparticles (GNPs) [20] would provide a stable and targeted delivery system for GDNF, thereby preserving and increasing the quantity of NMJs following VML. Specifically, the electrostatic coating of GDNF on the GNPs and their subsequent encapsulation within the BSGs will improve the bioavailability of the growth factor by increasing the amount that can be carried and released at the target site and by ensuring the stability and protection of the electrostatically bound growth factor. We further propose that exogenous GDNF can play crucial roles during different phases of injury. In the acute phase, it can augment existing levels and counteract degradation caused by inflammation [21, 22]. In the chronic phase, it can provide sustained support for reinnervation and recovery, particularly when endogenous GDNF levels decline [19].

## 2. Materials and Methods

### 2.1 Biosponge (BSG) Preparation

Sterile deionized (DI) water was heated to 50°C to make a 3 wt% Type A porcine skin gelatin (G2500-100G, Sigma Aldrich) solution. After complete dissolution, 20 mM of 1-ethyl-3-(3-dimethyl aminopropyl) carbodiimide (EDC; 22980, ThermoFisher) and 8 mM of n-hydroxysuccinimide (NHS; 130672-100G, Sigma Aldrich) were added to the gelatin solution. Rat tail collagen I (9.50 mg/mL; 354249, BD Corning) was diluted in 1X phosphate-buffered saline (PBS; 10010-023, Gibco) to a concentration of 3 mg/mL. In a 6-well plate, 3.5 mL of the gelatin solution was combined with 1.5 mL of the collagen solution and 42 μL of laminin (LM)-111 solution (6 mg/mL; 3446-005-01, R&D Systems). The final concentrations of the components were 21 mg/mL of gelatin, 0.9 mg/mL of collagen, 50 μg/mL of LM-111, 14.6 mM of EDC, and 5.6 mM of NHS. The solution was polymerized in the refrigerator (4°C) for 1 hour and was frozen overnight at -8°C. The well plate was then moved to -80°C for at least 72 hours, then lyophilized for 12-14 hours. After lyophilization, the BSGs were frozen at -20°C until needed.

### 2.2 Gelatin Nanoparticle (GNP) Synthesis

To synthesize negatively charged GNPs, a previously described desolvation method was followed [20]. Briefly, 80 mg of type B bovine skin gelatin (G9382-100G, Sigma) was treated with sodium hydroxide (0.1 N) until a pH of 10 was reached. This alkaline Type B gelatin solution was added dropwise using a syringe with a 21G needle to a solution containing 6.3 mL of acetone, 4.2 mL of autoclaved DI water, and 4.5 mL of poloxamer-188 (P556, Sigma; 6230A, Mirus), under mechanical agitation. Diisopropylcarbodimiide (DIC; A19292.30, ThermoFisher Scientific), a hydrophobic zero-length crosslinker, was added at a final concentration of 9.2 mg/mL to the gelatin solution. This solution was vortexed and then allowed to stir for 24 hours at room temperature for complete crosslinking to occur. The alkaline Type B GNPs were then isolated by centrifugation at 5000 RPM for 10 minutes. This resulted in a phase separation, allowing GNPs to be collected from the top layer. The collected GNPs were then frozen, lyophilized, and stored at -20°C until needed.

### 2.3 GNP and GDNF Conjugation and Characterization

The lyophilized GNP powder was weighed and added to sterile nuclease-free water to create a 1.0 mg/mL solution. BSG scaffolds were flash-frozen in liquid nitrogen and pulverized to obtain a powder, which was weighed and resuspended at a final concentration of 1.0 mg/mL. These solutions were sonicated (Fisherbrand Ultrasonic Processor, 500W, 20kHz, 40% amplitude, 1 minute. 5 sec. ON/5 sec. OFF pulse) on ice to prevent particle aggregation and potential heat-induced degradation. This process was repeated to obtain suspensions of both acidic GNP (GNP^+^) [20] and alkaline GNP (GNP^-^). The GNP and BSG suspensions were analyzed by DLS using a Zetasizer Nano ZS (Malvern Instruments) to determine their hydrodynamic size and zeta potential. At least three replicate measurements were taken per sample, as described previously [20]. In a separate experiment, GNP powder with or without murine GDNF (30 kDa; 450-40-50UG, PeproTech) was combined with potassium bromide in a mass ratio of 1:4, respectively. This mixed powder was compressed into a thin pellet and analyzed using Fourier-Transform Infrared Spectroscopy (FTIR) equipment (Cary 630, Agilent Technologies).

### 2.4 GDNF Encapsulation within Biosponges (BSG)

Alkaline type B GNPs were suspended at a concentration of 5.0 mg/mL in nuclease-free water. This solution was combined with murine GDNF (30 kDa; 450-40-50UG, PeproTech) to achieve final concentrations of 1.0 mg/mL GNP and 1 µg/mL GDNF, respectively. This GNP^-^ :GDNF solution was allowed to incubate at room temperature for 10 minutes for electrostatic interactions to occur.

BSG scaffolds were disinfected with 70% ethanol for 10 mins and then biopsied to a diameter of 6 mm to fit into a 24-well plate. The BSGs were then sterilized in the EcoVs 59S UVC LED sterilizing chamber for two cycles of 3 minutes and then rinsed in 1x PBS twice for 5 mins each. The BSG scaffolds were allowed to dry for 48 hours in a sealed desiccator at 4°C. The BSG scaffolds (6 mm disk) were then rehydrated individually in a GNP^-^: GDNF solution (100 µL) in the 24-well plate for 2 hours at 4°C.

### 2.5 Release of GDNF from Biosponges

After BSG rehydration with GNP^-^: GDNF solution, the BSG scaffolds were moved to a new 24 well-plate. Sterile 1x PBS (700 µL) was added to each well, and the well plate was incubated at 37°C. The PBS (700 µL releasates) was collected and replaced every 24 hours for the next seven days. Releasates and BSG scaffolds were frozen at –20°C until needed. The Mouse GDNF ELISA Kit (EEL117, Invitrogen) was used to quantify GDNF concentrations in the releasates. After 6 days of release, the biosponges were minced with scissors in a solution containing collagenase type II (1%), dispase (2.4 U/mL), calcium chloride (2.5 mM), and penicillin-streptomycin (1%) for 90 minutes at 37°C. The samples were vortexed every 15 minutes to ensure thorough digestion. The samples were centrifuged at 10,000 g, and the supernatants were collected for SDS-PAGE and western blotting analysis as described previously [23]. Briefly, the protein concentration was determined using the Pierce ^TM^ BCA Protein Assay kit. Equal amounts of reduced and denatured protein (50 µL) were resolved by SDS-polyacrylamide gel electrophoresis (SDS-PAGE) using 4–20% gels (Bio-Rad) and transferred onto nitrocellulose membranes. Equal protein loading was verified by Ponceau S staining. The membranes were probed using anti-GDNF polyclonal antibody (PAI-9524, Invitrogen, 1:250), HRP-conjugated secondary, and Clarity Western ECL Blotting Substrate. The membranes were imaged using the ChemiDoc Imaging System (Bio-Rad).

### 2.6 Release of GNP^-^:GDNF Conjugates and Their Temporal Effects on NSC-34 Neurite Growth

Previously developed BSGs were disinfected by rinsing in 70% ethanol and then biopsied to a diameter of 6 mm to fit into a 24-well Transwell® plate (Costar, REF 3413) with a 0.4 µm pore polycarbonate membrane insert. The BSGs were then sterilized in the EcoVs 59S UVC LED sterilizing chamber for two cycles of 3 minutes and then rinsed in 1x PBS twice for 5 mins each. They were then stored and allowed to dry for 96 hours at 4°C. Mouse Motor Neuron-Like Hybrid Cell Line (NSC-34; CLU140, Cedarlane Cellutions Biosystems Inc.) were seeded in the bottom chamber of a 24-well Transwell® plate at a density of 10^4^ cells per well (n=4/group) in complete proliferation media (DMEM-F12 + 10% FBS + 1% P/S). The cells were imaged after 24 hours and the media was replaced with complete differentiation media containing DMEM-F12, 1% FBS, 1% P/S, 1% MEM-Non-essential amino acid (NEAA), and 1 µM retinoic acid.

After 24 hours of culture, dry BSGs were separated into four groups: blank, GNP^-^-only, GDNF-only, and GNP^-^:GDNF and rehydrated as follows for 1 hour at 4 °C. The blank group BSGs were each rehydrated with 100 µL of sterile nuclease-free water. The GNP^-^-only BSGs were rehydrated with 100 µL of 5 mg/mL GNP^-^ suspension. GDNF-only BSGs were rehydrated 100 µL of 1 µg/mL GDNF suspension in sterile nuclease-free water. The GNP^-^:GDNF BSGs were rehydrated with 100 µL of a solution that included 5 mg/mL GNP with 1 µg/mL GDNF, which had been incubated at room temperature for 10 minutes to facilitate electrostatic conjugation. BSGs were placed in transwell inserts and placed on top of the wells containing NSC-34 cells. On days 3, 5, and 7, transwell inserts were removed from the wells and stored under aseptic conditions in the biosafety cabinet, and the cells were reimaged. Following imaging, differentiation media was replaced, and the transwell inserts were returned to their original wells. An XTT assay was performed after imaging on day 7. Images showed NSC-34 neurite extensions, and their length was manually quantified using ImageJ (or Fiji; National Institutes of Health).

### 2.9 Animal Experiments

All animal work was conducted in compliance with the Animal Welfare Act, the implemented Animal Welfare Regulations, and in accordance with the principles of the Guide for the Care and Use of Laboratory. All animal procedures were approved by Saint Louis University’s Institutional Animal Care and Use Committee (Animal Protocol # 2645).

Male Lewis rats (10-12 weeks old; Charles River Laboratories) were housed in a vivarium accredited by the Association for Assessment and Accreditation of Laboratory Animal Care International, which provided water and food *ad libitum*. The rats were randomly assigned to experimental groups. The animals were weighed and anesthetized using 2.5% isoflurane. The surgical site was shaved and aseptically prepared using 10% povidone–iodine and 70% isopropyl alcohol. The skin of the lower leg was laterally incised to reveal the tibialis anterior (TA) muscle. The skin was separated from the fascia and underlying musculature using blunt dissection. An incision was made on the fascia, and it was gently separated from the TA muscle using a blunt probe. The TA muscle was bluntly separated from the extensor digitorum longus (EDL) and the tibia. To create the VML injury, a flat metal spatula was inserted underneath the TA muscle. A 6-mm biopsy punch was used to remove ∼20% of the TA muscle’s mass. The biopsied mass was weighed for consistency. Any bleeding was controlled with light pressure using a sterile gauze.

The injury was treated with 6 mm disks of either blank GNP BSG or with GNP+GDNF-loaded BSG (n=4/group). The surgery was performed bilaterally, with both legs receiving the same treatment (i.e., n=8 muscles/group). The metal spatula was removed after the BSG (6-mm diameter) was placed in the defect. The fascia was pulled over the BSG and sutured together with 6-0 Prolene sutures (Ethicon). The skin incision was closed using simple interrupted Prolene (5-0; Ethicon) sutures. Sustained-release buprenorphine (1mg/kg) was administered subcutaneously at the nape of the neck for pain management. Investigators were blinded to group allocation during data collection, and animals were identified by a numerical code.

After 6 weeks of recovery, all animals underwent peak isometric torque measurement followed by euthanasia. The TA muscle was then harvested, weighed, and pinned in Sylgard^TM^ (24236-10, Electron Microscopy Sciences) dishes to retain its shape. The pinned muscles were then fixed in 10% formalin for an hour. The pins were then removed, and the muscle was transferred to a 6-well plate. The muscle was submerged in 5% sucrose solution, then in 10% sucrose solution, for 30 minutes each at room temperature. It was then incubated in a 20% sucrose solution overnight at 4°C [24]. After overnight incubation, TA muscles were embedded in 25 mm x 25 mm x 5 mm base molds using optimal cutting temperature (OCT) compound (23-730-571, Fisher Healthcare). The OCT molds were preserved for histological analysis using 2-methylbutane (Fisher Scientific), supercooled in liquid nitrogen to approximately –95°C.

### 2.10 Peak Isometric Torque Measurement

*In* vivo function testing of the anterior crural muscles was performed 6-weeks post-injury, as described previously [10, 12, 23, 25–28]. The rats were anesthetized using isoflurane (1.5-2.0%) on a heated pad. The skin was shaved, and the animals were moved to a heated test apparatus (Model 806D, Aurora Scientific) to maintain normal body temperature. The knee was then clamped to secure the leg. After ensuring the leg was secured, the animal’s foot was positioned on the footplate of the dual-mode motor (Model 305C, Aurora Scientific). The heel was placed in the groove of the footplate, and adhesive tape was used to fasten the foot to the pedal. The foot was positioned perpendicularly to the tibia, forming a 90° angle. Two platinum electrodes were inserted subcutaneously on each side of the common peroneal nerve to stimulate the anterior crural muscles and to elicit dorsiflexion. Optimal current amplitude (30-35 mA) was set with a series of twitches. Isometric tetanic contractions were elicited at 150 Hz (0.1 ms pulse width, 300 ms train). The data was acquired using the Dynamic Muscle Control/Dynamic Muscle Analysis (DMC/DMA) software (Aurora Scientific).

### 2.11 Histological Analysis

The frozen TA muscles embedded in OCT were mounted face up on cryostat chucks using OCT compound and cryosectioned using a Leica CM1850 cryostat to obtain longitudinal cross-sections (60 µm). Sections were mounted on Superfrost^TM^ Plus Gold Slides (22-035813, Fisher Brand).

Longitudinal sections (60 µm) taken at a depth of 720-960 µm from the muscle’s surface were fixed in 200-proof methanol chilled to –8 °C for 15 minutes, followed by permeabilization in 0.1% Triton X-100 for 15 minutes. Blocking was performed using 3% BSA, 0.05% Triton X-100, and 5% goat serum for 1 hour. The NMJs were identified by using antibodies against acetylcholine receptors [Alpha-bungarotoxin (α-BTX) conjugated with Alexa Fluor 488, 1:50; B13422, Invitrogen], pre-synaptic vesicles [Synaptophysin (Syn) monoclonal antibody, 1:100; MA5-14532, ThermoFisher], and axons [Anti-Neurofilament (NF) H antibody, 1:300; AB5539, MilliporeSigma][6]. Antibodies for α -BTX, Syn, and NF were utilized to label the following structures, i.e., post-synaptic endplate, pre-synaptic endplate, and axons, respectively. Following blocking, the sections were incubated overnight at 4 °C with Syn and NF antibodies diluted in incubation buffer (3% BSA, 0.05% Triton X-100). The following day, sections were incubated with appropriate fluorochrome-conjugated secondary antibodies (Alexa Fluor 594, 1:500; A11042 and A11012; Invitrogen) along with α-BTX antibody in incubation buffer for 1 hour at room temperature. Sections were washed in PBS (3x for 5 minutes each), followed by incubation in ProLong^TM^ Gold Antifade Mountant with DNA stain DAPI (P36931, Molecular Probes by Life Technologies) to identify cell nuclei. The stained slides were mounted with a coverslip and sealed with nail polish. Slides were imaged no later than 24 hours post-sealing.

A separate set of longitudinal sections (60 µm) taken at a depth of 960-1200 µm from the muscle’s surface was stained with hematoxylin and eosin (H&E) to visualize myofiber morphology and BSG remodeling. Using the freehand selection tool in ImageJ, the BSG area was manually quantified from composite H&E images.

### 2.12 NMJ analysis

A total of 1055 individual NMJs were imaged and analyzed for this study, with an average of 44±25 *en face* NMJs analyzed per muscle sample. Imaging was performed on the Andor Dragonfly 200 spinning disk confocal (Oxford Instruments), mounted on a Leica DMi8 base, equipped with a HC Fluotar 25x/0.95 W VISIR water-immersion lens.

Qualitative analysis of NMJs in confocal images was performed manually. NMJs were classified by the extent of overlap between the pre-synaptic nerve terminal stained with Syn+NF and post-synaptic acetylcholine receptors stained with α-BTX [29]. Complete overlap between pre- and post-synaptic terminals was defined as >75% colocalization between α-BTX^+^ and Syn+NF^+^ structures. Partial overlap was defined as 25-75% colocalization between α-BTX^+^ and Syn+NF^+^ staining. Finally, no overlap was defined as <25% colocalization between α-BTX^+^ and Syn+NF^+^ staining.

Quantitative analysis was performed by two blinded investigators using a modified version of the NMJ-Morph methodology [30] to quantify the following: presynaptic area, postsynaptic area, colocalization area, and number of fragments. Image stacks of at least 60 optical slices (1 μm step size) comprising three channels (16-bit RGB) were used for analysis of pre- and postsynaptic NMJ structures. Using Fiji, the 16-bit image stacks were separated into 8-bit maximum intensity projections. The image intensity threshold was determined manually, with visual confirmation, and the corresponding threshold values were recorded. To manually set the threshold, two maximum intensity images were opened, using one as a reference; the threshold was adjusted to match the reference, and the threshold value was recorded. Using the ‘polygon’ tool and the ‘measure’ function, the 2-D planar area of pre- and post-synaptic structures was determined. Previously recorded threshold values were used in the ‘colocalization’ plugin to determine the area of overlap between pre- and post-synaptic structures. This was used to calculate the area of colocalization as a percentage of the post-synaptic area. Finally, using the ‘analyze particles’ function in Fiji, the total number of motor endplate fragments was obtained (**Supp. Fig. 1**). Following quantification, the four measurements, i.e., Presynaptic area, postsynaptic area, percent colocalization, and number of fragments, were categorized into size bins to obtain the size distribution frequency.

### 2.13 Atomic Force Microscopy (AFM) analysis

A high-performance MFP-3D-Bio atomic force microscope (AFM; Oxford Instruments) was used to analyze the mechanical properties of muscle tissues. A set of longitudinal sections (60 µm) taken at a depth of 480-720 µm from the muscle’s surface were used for analysis. Tip-less AFM cantilevers (ARROW-TL1Au, nominal spring constant: 0.03 N/m) were modified by fixing a 25-μm polystyrene bead, as described earlier [31]. At least 25 indentation curves were obtained over a 100 × 100 µm scan area at various locations on the tissues, with a setpoint of 5 nN and a constant scan rate of 0.15 Hz. The Hertzian model was used to calculate Young’s modulus (kPa). The data were classified into three distinct regions of the injured muscle: the injury or defect zone, the injury adjacent zone, and the distant zone.

### 2.14 Statistical Analysis

Data are presented as a mean ± standard error of the mean. Unpaired t-tests and one-way analysis of variance (ANOVA) were used to determine significant differences between groups. Two-way ANOVA was used to determine if there was a significant interaction or main effect between variables. A Fisher’s LSD post-hoc comparison was utilized to identify the source of significance with p < 0.05 unless otherwise specified in figure captions. GraphPad Prism 10 for Windows was used to perform statistical analyses and graph data.

## 3. Results

### 3.1 GDNF release kinetics

Zeta potential measurements confirmed that alkaline Type B GNPs were negatively charged (-12 mV) and Type A biosponges were positively charged (11.8 mV). GDNF-coated GNPs showed a reduced charge (-8.6 mV) likely due to the coating of GDNF on the GNP surface (**Fig. 1A**). However, all three groups were statistically different from each other (one-way ANOVA, p<0.0001). The GDNF release study demonstrated a gradual, consistent decline in concentration over 6 days *in vitro* (**Fig. 1B**). A third-order (cubic) polynomial fit was used for the release data (R^2^ = 0.3334). Western blot results indicated that GDNF remained within the BSG structure even after 6 days of release (**Fig. 1C**).

**Fig.1.**
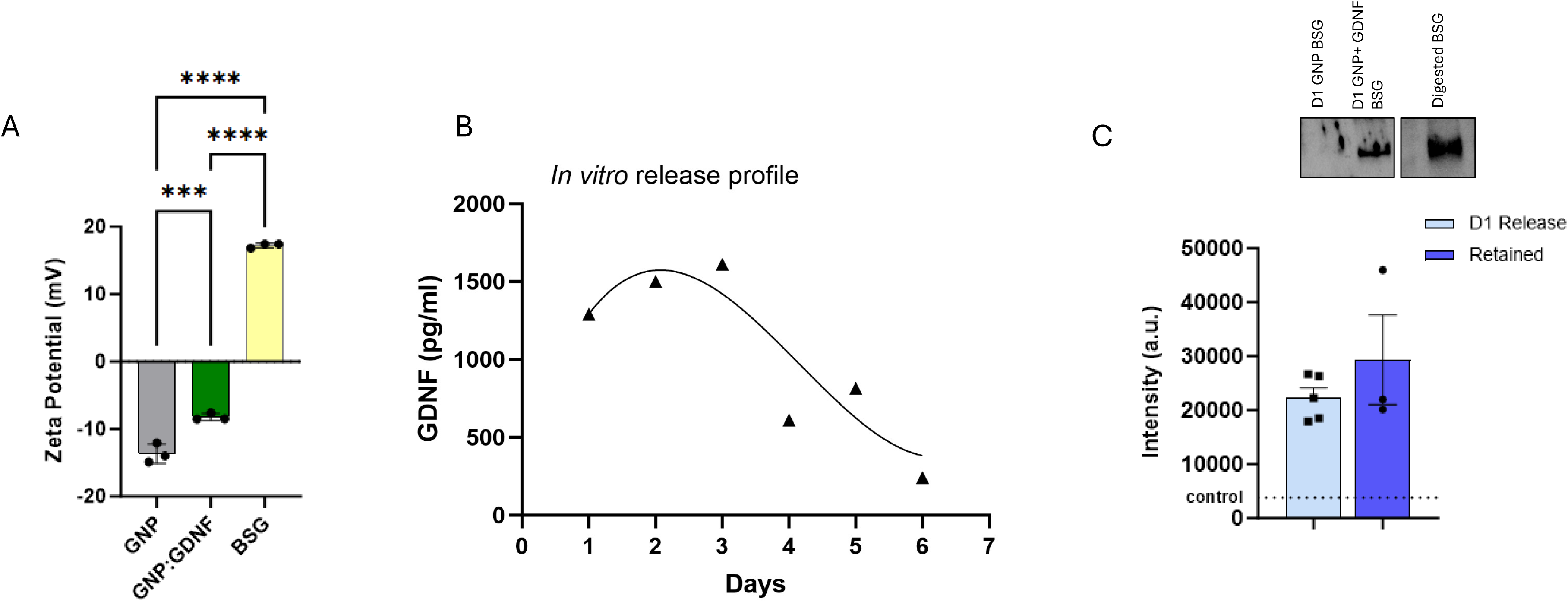
(A) The zeta potential was measured for GNP, GNP:GDNF conjugates, and the biosponges (BSG). Data analyzed using one-way ANOVA. “*” indicates a significant difference (*p<0.05, ***p<0.001, ****p<0.0001) between groups. (B) The *in vitro* release of GDNF from BSGs was quantified using ELISA. (C) The amount of GDNF retained within BSGs after 6 days of *in vitro* release study was measured following BSG digestion using western blotting.

FTIR spectra of both acidic **(Fig. 2A)** and alkaline GNPs **(Fig. 2C)** demonstrate increases in transmittance upon conjugation with GDNF **(Fig. 2B-D**). These changes occur primarily around the Amide I (1600-1700 cm⁻¹) and Amide II (1510-1580 cm⁻¹) bands. Sharper amide bands typically suggest a reduction in conformational flexibility [32–36]. These changes indicate that the interaction of GDNF with GNP has induced a more ordered or specific structural arrangement within the conjugate. The structure of GDNF is known to feature both positive and negative residues on its surface [37], so differences in spectral intensity between GDNF conjugated to positively and negatively charged GNPs are expected.

**Fig.2.**
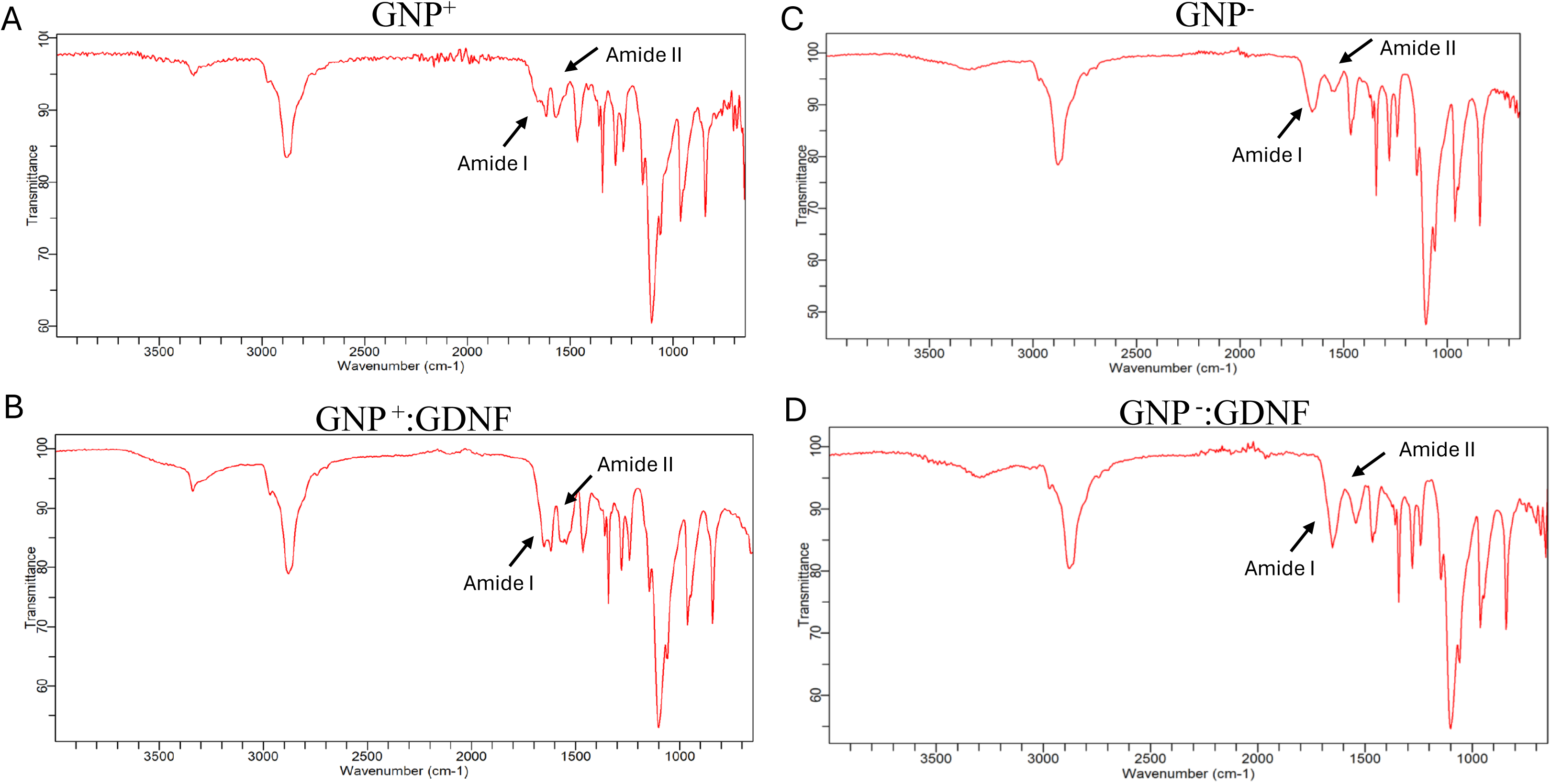
The FTIR spectra for (A) positively charged GNPs, and (B) GNP^+^:GDNF conjugates as well as for (C) negatively charged GNPs, and (D) GNP^-^:GDNF conjugates. The addition of GDNF led to sharper and more enhanced Amide I and II bands in the gelatin nanoparticle spectra suggesting possible interaction between gelatin and GDNF and indicating the electrostatic conjugation of GDNF to the gelatin nanoparticles.

### 3.2 Bioactivity of GDNF on NSC-34 cells

NSC-34 cellular morphology on days 1, 3, 5, and 7 of culture is shown in **Fig. 3A**. Neurite extension quantification on days 3 and 5 showed that the groups containing GDNF exhibited significantly longer neurite extensions as compared to the groups without GDNF (**Fig. 3B)** (2-way ANOVA, Interaction p=0.7525, Time effect, p=0.0347, Treatment effect, p<0.0001). Only on day 3, the GNP^-^: GDNF conjugate group resulted in significantly longer neurites than the GDNF only group (p=0.0259). Quantifiable neurites were not observed on day 1, and cellular confluency obscured measurements on day 7. Therefore, neurite extensions were not quantified on days 1 and 7. A size distribution analysis of neurite length was also performed. Quantification of neurite extensions on day 3 shows a significant increase in the percentage of neurites greater than 30 µm in the GNP^-^: GDNF conjugate group as compared to all other groups (**Fig. 4A-B**; 2-way ANOVA, Interaction, p<0.0001, Size factor, p<0.0001, Treatment factor, p>0.9999). Notably, the percentage of neurite extensions within the > 30 µm size bin increased from ∼38% on day 3 (**Fig. 4A**) to ∼60% on day 5 (**Fig. 4B).** These results indicate that GNP: GDNF conjugates result in significantly longer neurite extensions compared to unconjugated GDNF.

**Fig.3.**
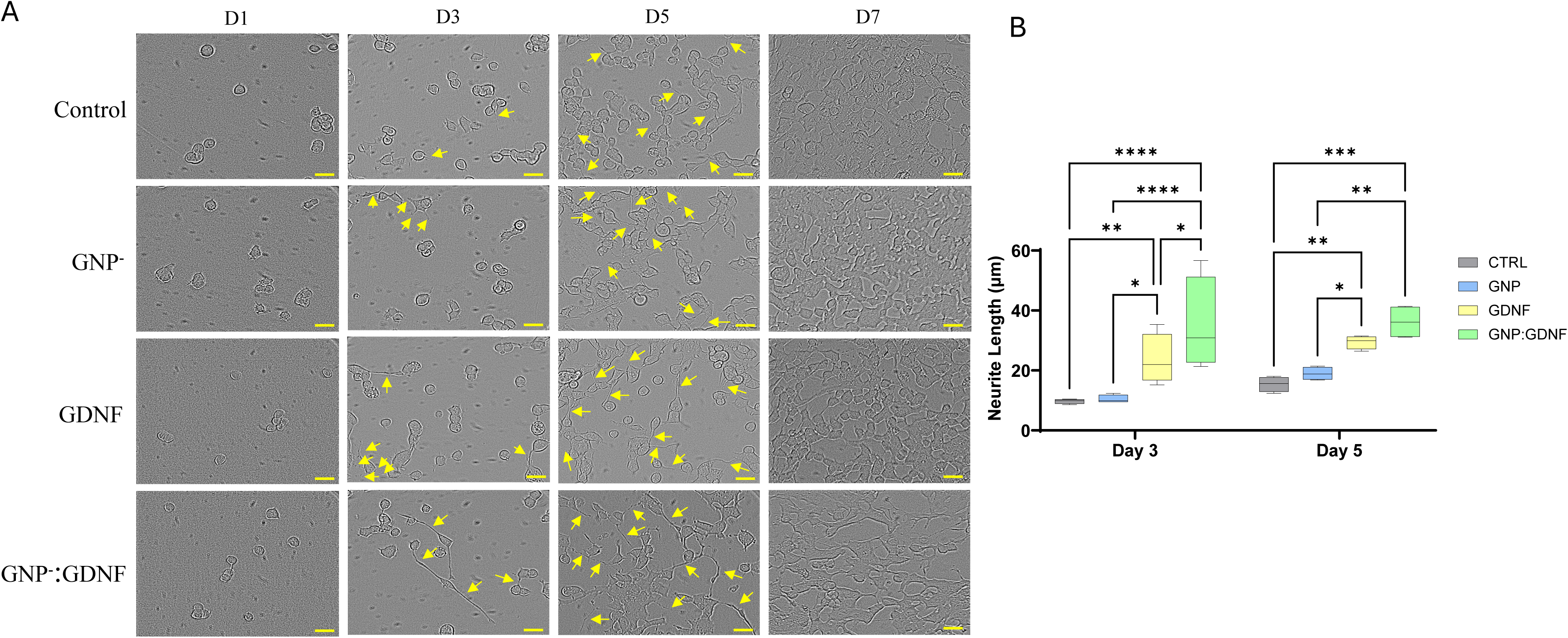
(A) NSC-34 motor neuron cells were cultured in transwell setups that subjected the cells to four different types of biosponges (BSG), i.e., Control (CTRL), GNP, GDNF, and GNP+GDNF. (B) Neurite extensions were quantified on days 3 and 5. The GDNF-containing groups showed significantly longer neurite extensions than the groups without GDNF. The data were analyzed using a 2-way ANOVA. “*” indicates a significant difference (*p<0.05, ***p<0.001, ****p<0.0001) between groups.

**Fig.4.**
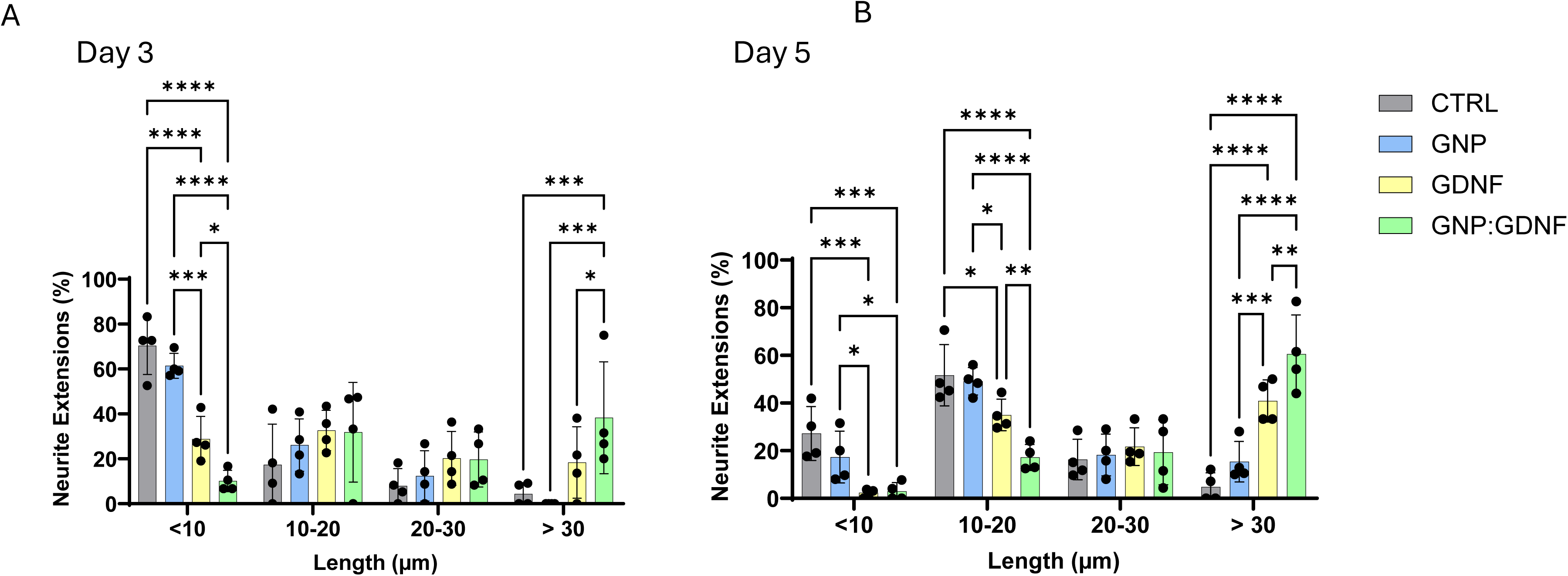
The size distribution analysis of neurite extensions from NSC-34 cells showed that GNP+GDNF containing biosponges resulted in significantly longer neurite extensions in the >30 μm size range on days 3 and 5. The data was analyzed using 2-way ANOVA. “*” indicates a significant difference (*p<0.05, ***p<0.001, ****p<0.0001) between groups.

### 3.3 Muscle Mass and Function

The biopsy mass removed to create the VML injury was consistent between the experimental groups (**Fig. 5A)** (t-test, p = 0.9074). The TA muscle mass was measured at 6 weeks post-injury. Post-normalization of muscle mass to body weight of the animal, the results showed ∼33.5% (GNP) and ∼26.5% (GNP+GDNF) deficits relative to the uninjured muscle. However, no differences between treatment groups were observed (**Fig. 5B**) (One-way ANOVA, p<0.0001; U vs. GNP, p<0.0001; U vs. GNP+GDNF, p =0.0002).

**Fig. 5.**
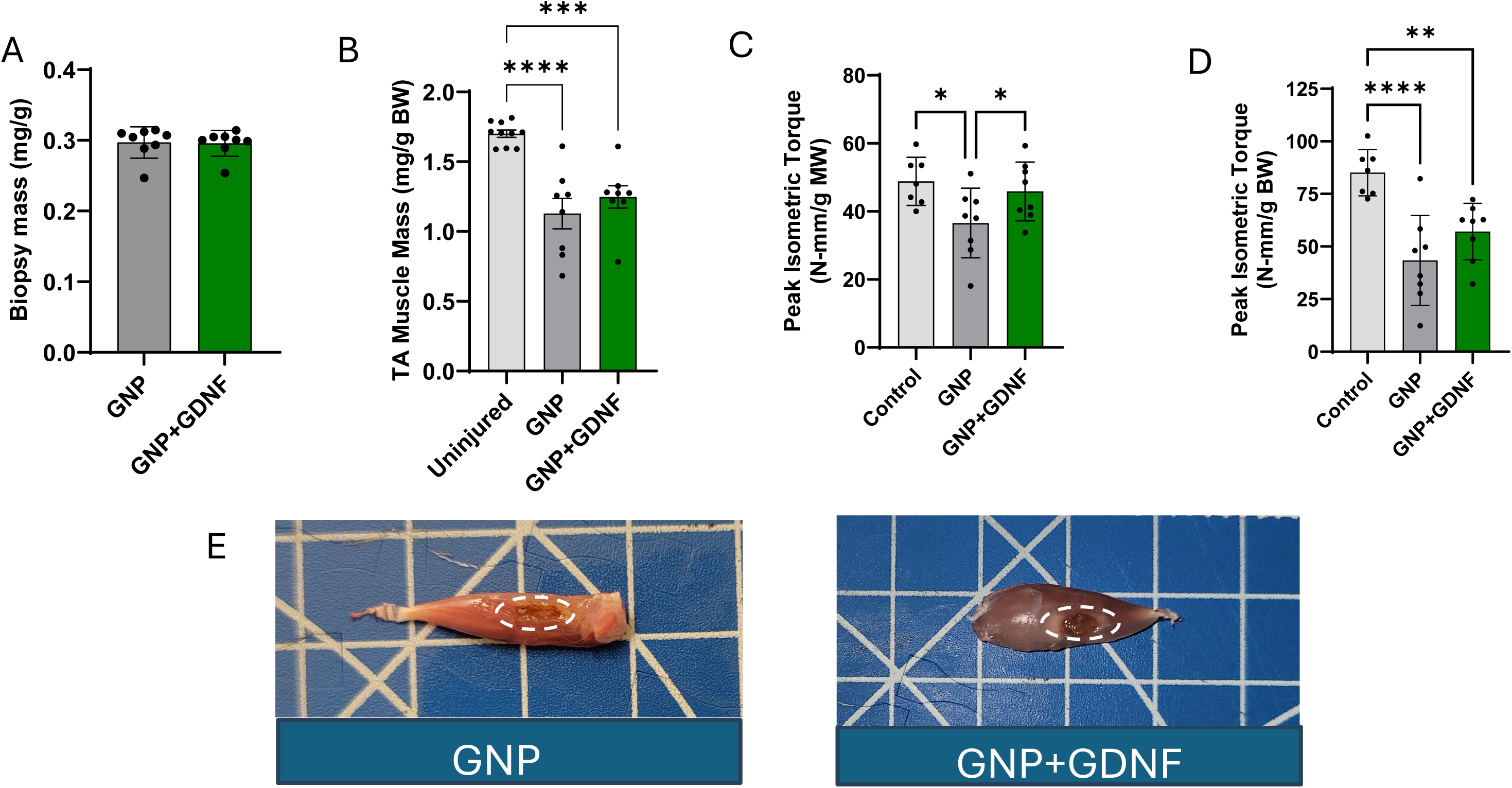
(A) The biopsy mass removed to create the VML defect was measured to ensure consistency in the procedure. (B) The TA muscle’s mass was normalized to the body weight of the animal. Peak isometric torque measurements were performed and normalized to the (C) mass of the TA muscle as well as the (D) body weight of the animal (E) Photographs of the excised TA muscle are presented. In both groups, a remodeled biosponge, outlined in white dashes, is visible. The data was analyzed using one-way ANOVA. “*” indicates a significant difference (*p<0.05, ***p<0.001, ****p<0.0001) between groups.

Peak isometric torque was measured at 6 weeks post-injury and showed a significant increase (∼25%) with GNP+GDNF treatment relative to the GNP only group, when normalized to the TA muscle mass (**Fig. 5C**) (one-way ANOVA, p = 0.0337). However, no differences were found when comparing the peak isometric torque normalized to the body weight of the animal between GNP only and GNP+GDNF (**Fig. 5D**; One-way ANOVA, p=0.0003). The BSG could be visually identified in the excised TA muscles in both groups, as shown in the photographs (**Fig. 5E**).

### 3.4 Biosponge Remodeling and AFM Analysis

The BSG area quantified from H&E-stained muscle histological sections is shown in **Fig. 6A**. The GNP+GDNF group showed a higher area of BSG relative to that of the GNP only group, suggesting slower remodeling and degradation in the presence of GDNF (**Fig. 6B**; t-test, p=0.0376). AFM analysis revealed that the Young’s modulus of the VML-injured muscle (irrespective of treatment) is significantly lower than that of the uninjured controls **(Fig. 7A-B**; Two-way ANOVA, interaction, p=0.0193, regional factor, p<0.0001, treatment factor, p<0.0001). Additionally, the modulus varies regionally within the injured muscle, with the lowest stiffness in the injury and adjacent zones and higher values in distant areas.

**Fig. 6.**
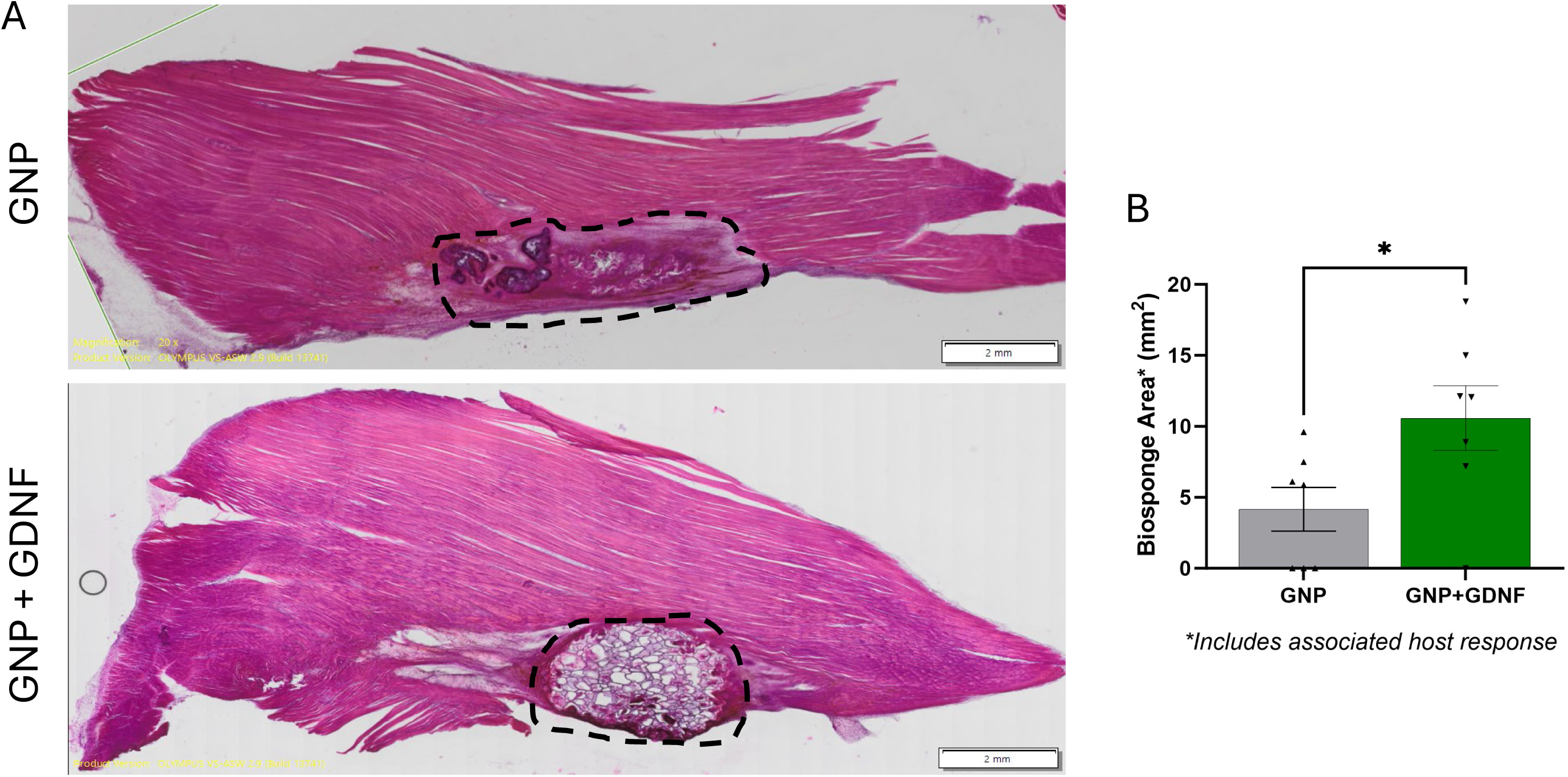
(A) Muscle longitudinal sections were stained with H&E at 6 weeks post-injury (scale bars = 2 mm). The embedded biosponge is outlined in black dashed lines. (B) Biosponge area was quantified manually, and its remodeling was slower when GDNF was incorporated within it. Data analyzed using unpaired t-test (*p<0.05).

**Fig.7.**
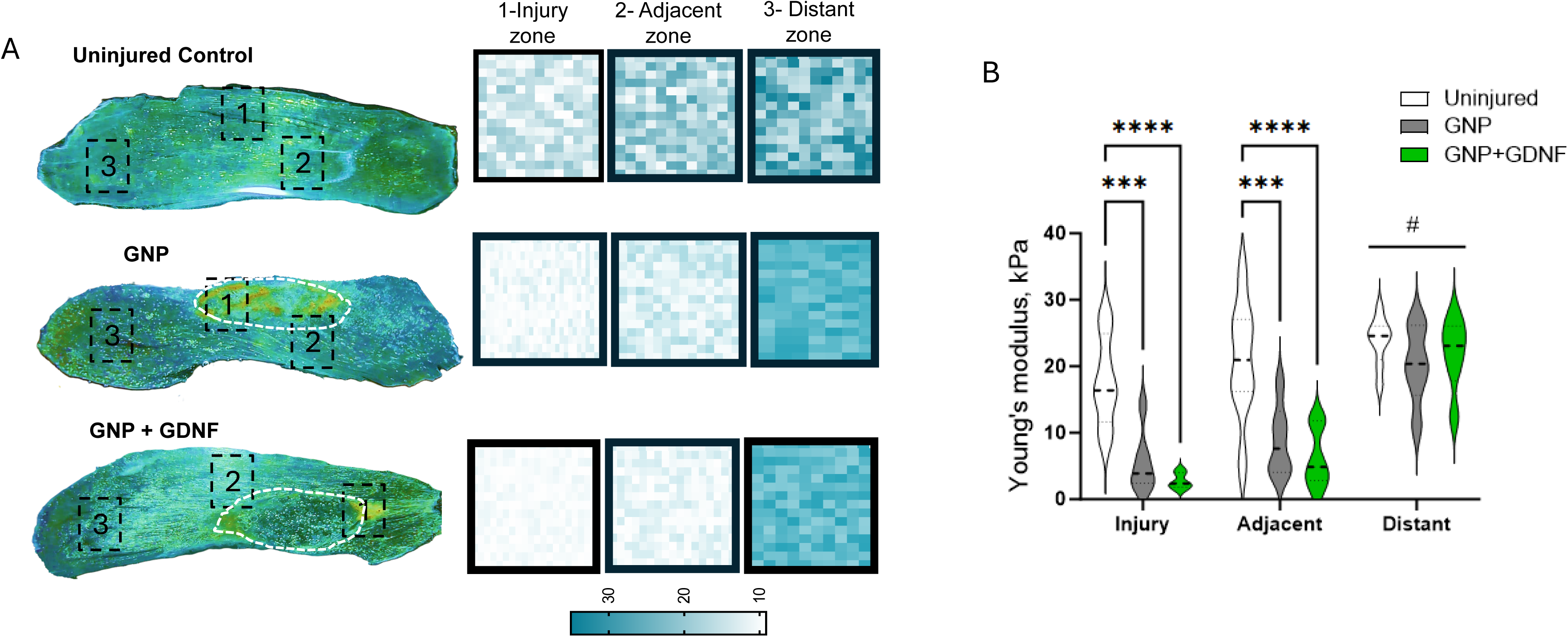
(A) The longitudinal sections imaged using the AFM setup were analyzed in injury (1), adjacent (2), and distant (3) regions. The force maps show the elastic modulus values of the muscle tissue in a color scale. (B) AFM analysis revealed a significant reduction in Young’s modulus in both injury groups (irrespective of treatment) relative to the uninjured control group, exclusively in the injury and adjacent regions. Data was analyzed using two-way ANOVA. “*” indicates a significant difference (*p<0.05, **p<0.01, ***p<0.001, ****p<0.0001) between groups.

### 3.5 Neuromuscular junction analysis

Representative confocal images for each experimental group showing the pre- and post-synaptic structures labeled with NF+SYN and α-BTX are presented in **Fig. 8A**. Semi-quantitative assessment of NMJs in the confocal images showed an average of ∼71% complete, 20% partial, and 14% no overlap in the uninjured TA muscles. The GNP+GDNF group exhibited 71% complete, 11% partial, and 18% no overlap between pre- and post-synaptic structures, compared to 42% complete, 40% partial, and 18% no overlap in the GNP group (**Fig. 8B**; Two-way ANOVA Interaction p= 0.0054, Treatment Factor p=0.9643, Overlap Factor p<0.0001). NMJs exhibited a pronounced improvement between pre- and post-synaptic structure overlap in the GNP+GDNF group as compared to the GNP-only group (Complete overlap, GNP vs. GNP+GDNF, p=0.0113).

**Fig. 8.**
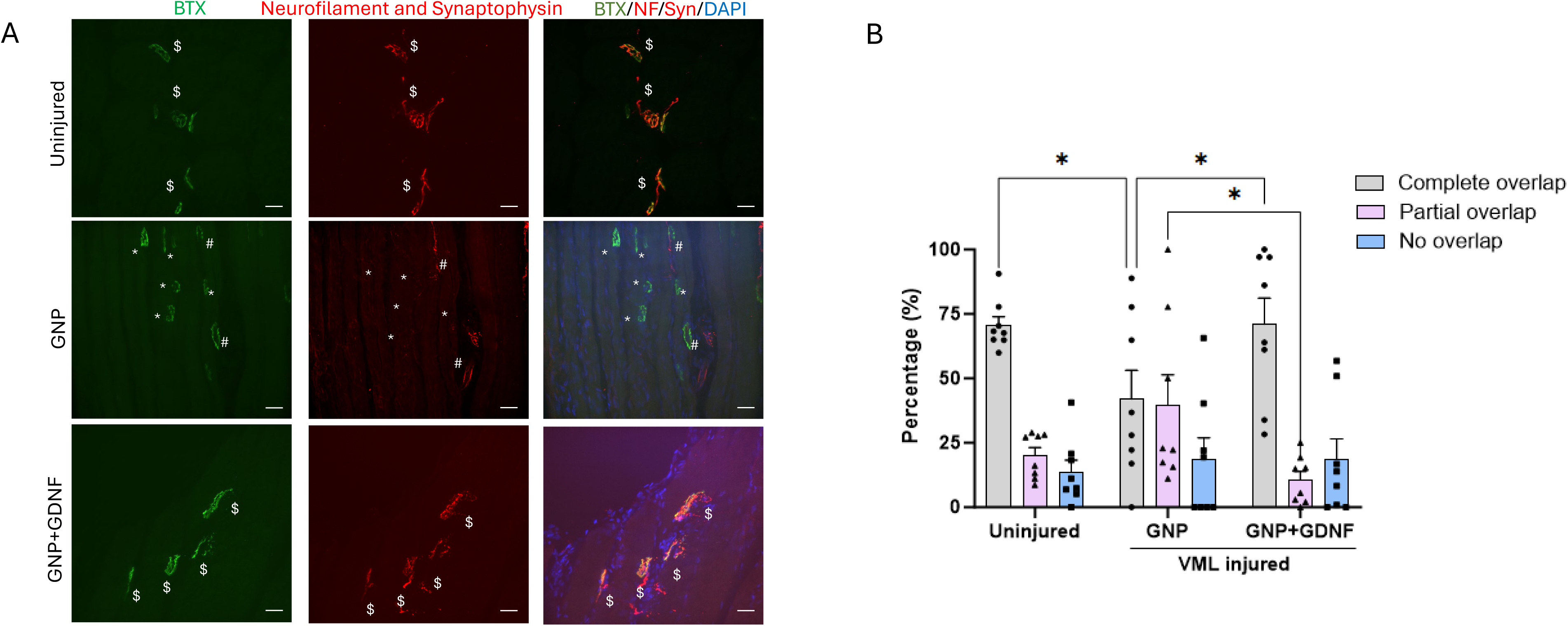
(A) Muscle longitudinal sections were stained with anti-neurofilament, anti-synaptophysin, and alpha-bungarotoxin to visualize pre- and post-synaptic structures 6 weeks post-injury (Scale bar = 20 µm). (B) Pre- and post-synaptic structures showing complete, partial, or no overlap were quantified manually. The annotations on the images indicate no overlap (*), partial overlap(#), and complete overlap($). Data was analyzed using 2-way ANOVA. On the bar graphs, “*” indicates a significant difference (*p<0.05, **p<0.01, ***p<0.001, ****p<0.0001) between groups.

Representative images in **Fig. 9A** depict both individual and merged, presynaptic (red) and postsynaptic (green) terminals, including the overlapping region (colocalization; white) for a fully innervated and a partially denervated neuromuscular junction (NMJ). A quantitative assessment of presynaptic and postsynaptic terminal colocalization revealed no significant difference in the overall mean percentage between the treatment groups. The mean percentage of colocalization was found to be ∼51% in the uninjured group, ∼34% in the GNP-treated group, and ∼49% in the GNP+GDNF-treated group **(Fig. 9B**). However, upon categorization into bins based on overlap percentage, the GNP group displayed a higher frequency of low colocalization (e.g., 10-30% overlap), in contrast to the GNP+GDNF group, which exhibited a predominant high colocalization (e.g., 70-90% overlap) **(Fig.9C**; Two-way ANOVA, Interaction p=0.0003, Colocalization factor p<0.0001, Treatment factor p>0.9999)

**Fig. 9.**
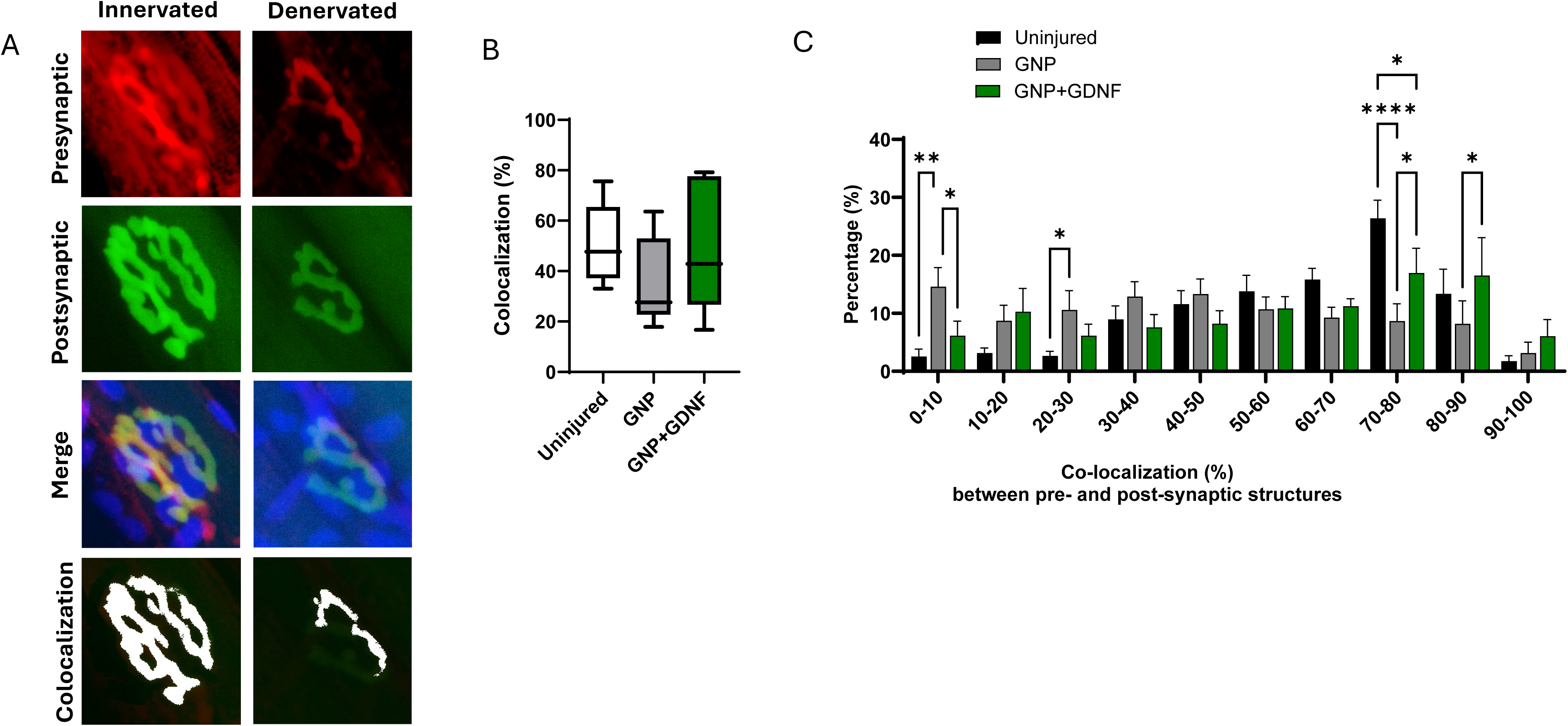
(A) Representative images show pre- and post-synaptic terminals individually and merged along with the colocalized regions (white) for an innervated and denervated NMJ. (B) Blinded investigators quantitatively assessed the extent of colocalization between the pre- and post-synaptic structures. The box-and-whiskers plot shows the minimum and maximum values, along with the median. (B). Binned analysis revealed that the GNP group had significantly higher amounts of 0-10% overlap, while the GDNF group had significantly higher amounts of 70-90% overlap. The data was analyzed using 2-way ANOVA. On the bar graphs, “*” indicates a significant difference (*p<0.05, **p<0.01, ***p<0.001, ****p<0.0001) between groups.

The post-synaptic area was measured to be 287 µm^2^ in the uninjured group, 388 µm^2^ in the GNP group, and 320 µm^2^ in the GNP+GDNF group, showing a 35% and 11.5% increase relative to the uninjured group, respectively **(Fig. 10A).** A significant difference was observed between the uninjured group and the GNP group (Kruskal-Wallis, p=0.0155). Distribution analysis also revealed a shift towards larger postsynaptic terminals, primarily in the GNP alone group **(Fig. 10B**).

**Fig. 10.**
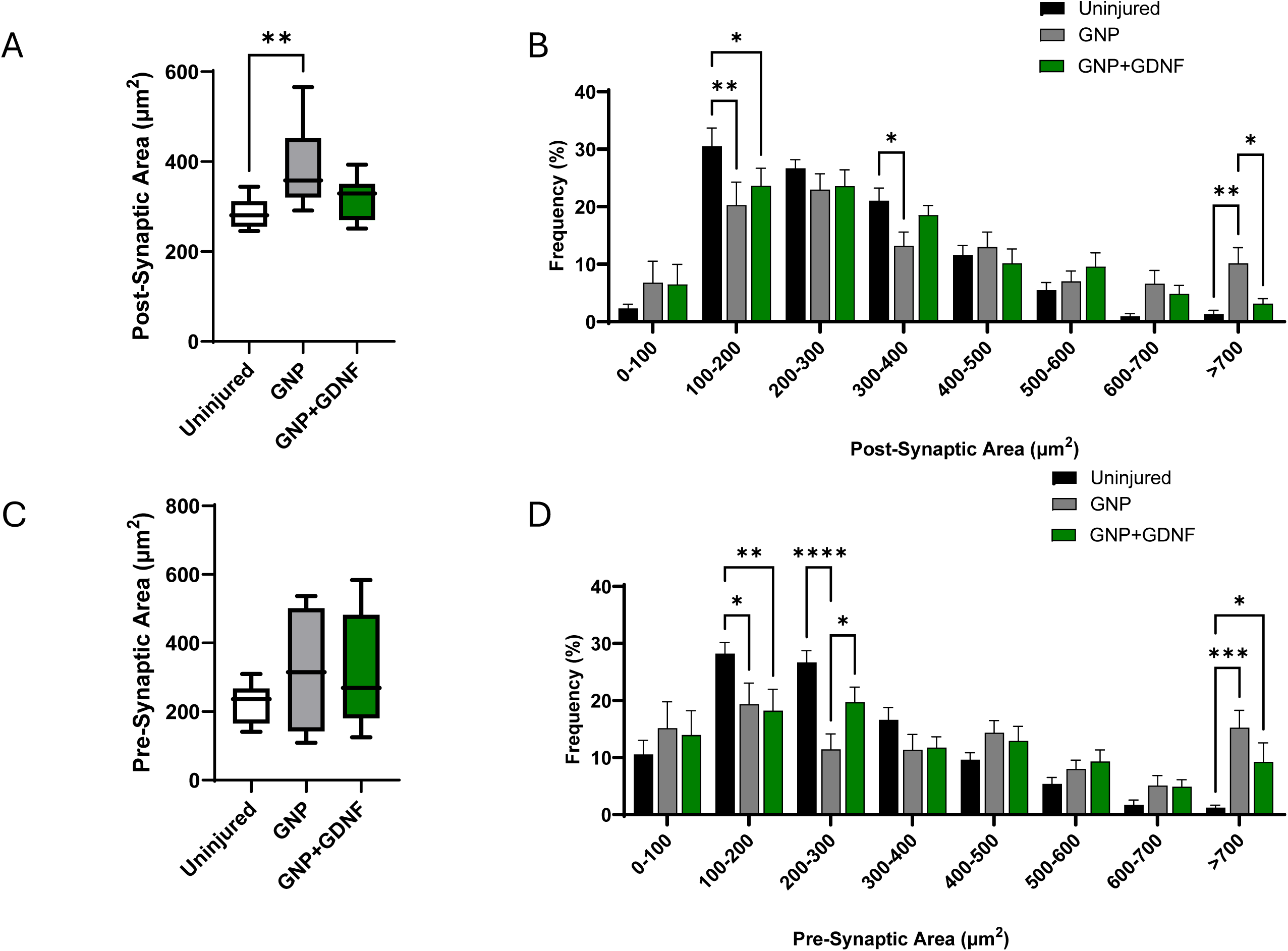
(A) The mean post-synaptic area was significantly higher in the GNP group relative to the uninjured control group (Kruskal-Wallis test). (B) Post-synaptic area distribution analysis reveals a shift towards larger postsynaptic terminals, primarily in the GNP alone group (2-way ANOVA). (C) The pre-synaptic area was similar between groups (Kruskal-Wallis test). (D) Pre-synaptic area distribution analysis shows a shift toward larger pre-synaptic terminals in both the GNP and GNP+GDNF groups (2-way ANOVA). On the bar graphs, “*” indicates a significant difference (*p<0.05, **p<0.01, ***p<0.001, ****p<0.0001) between groups. The box-and-whiskers plot shows the minimum and maximum values, along with the median.

The pre-synaptic area was measured to be 225 µm^2^ in the uninjured group, 320 µm^2^ in the GNP group, and 314 µm^2^ in the GNP + GDNF group, revealing a 42% and 40% increase relative to the uninjured group, respectively **(Fig. 10C**). However, statistical analysis showed no significant differences between the groups (Kruskal-Wallis, p=0.6151). However, distribution analysis revealed a shift toward larger pre-synaptic terminals in both the GNP and GNP+GDNF groups **(Fig. 10D).**

Representative images of fragmentation in motor end plates are depicted in **Fig. 11A**. Treatment of VML with BSG containing GNP-alone led to an increase in postsynaptic fragmentation **(Fig. 11B**; Kruskal-Wallis test, p=0.0229). However, when the BSG was supplemented with GNP+GDNF, the extent of fragmentation was comparable to that of uninjured control muscles. Size distribution analysis revealed an increase in the percentage of endplates with greater than four fragments in the GNP alone group relative to the uninjured muscles **(Fig. 11C**; Two-way ANOVA, Interaction p=0.0159, Fragmentation factor p<0.0001, Treatment Factor p>0.9999).

**Fig. 11.**
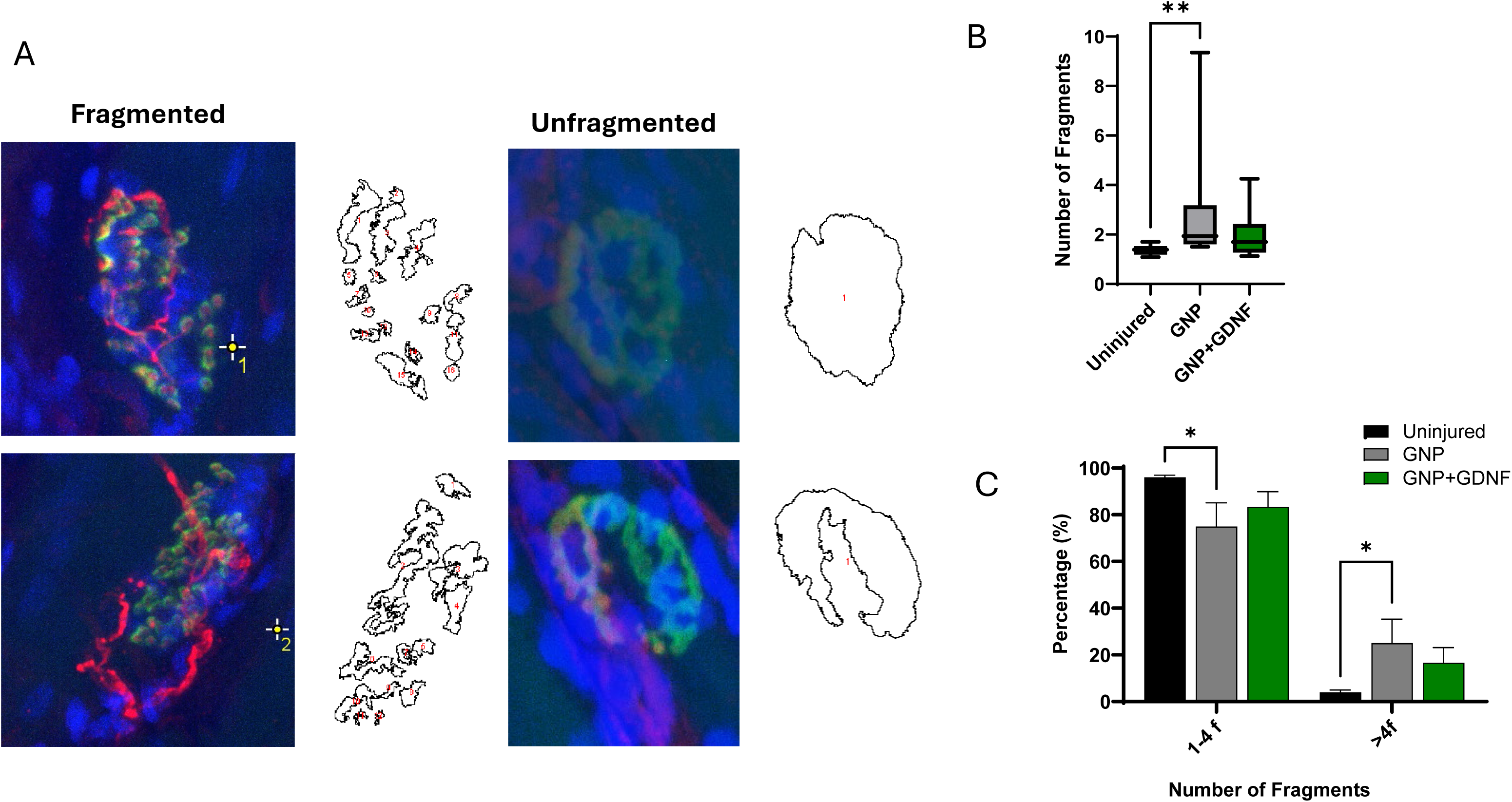
(A) Representative images of fragmented and unfragmented endplates are presented with particle analysis outputs from ImageJ. (B) The total number of fragments was quantified, and the data were analyzed using the Kruskal-Wallis test. (C) The degree of fragmentation of the motor endplate was quantified. Injured groups displayed higher amounts of >4 fragments as compared to the uninjured group. Data analyzed using 2-way ANOVA. On the bar graphs, “*” indicates a significant difference (*p<0.05, **p<0.01, ***p<0.001, ****p<0.0001) between groups. The box-and-whiskers plot shows the minimum and maximum values, along with the median.

## Discussion

The most salient finding of this study is that treatment with a BSG scaffold encapsulating GDNF improved the number of functional NMJs as well as peak isometric torque production in VML-injured muscles. These results suggest that the biosponges infused with GDNF are an effective strategy for enhancing muscle recovery after VML.

In a previous study, Greising *et al.* [6] observed a leftward shift in the presynaptic terminal size distribution, suggesting a decrease in size, coupled with a rightward shift in the postsynaptic terminal size distribution, indicating an increase in size following VML injury. A shrinking of the presynaptic terminal would suggest that ACh molecules are not being consistently released. The enlargement of postsynaptic terminals suggests a compensatory adaptation in which the AChRs attempt to maximize their chances of capturing ACh. Additionally, enhanced fragmentation of the postsynaptic terminal was also observed by Greising *et al.* [6]. A similar pathology has also been observed in dystrophic mice [8, 38].

In contrast, we observed that both the presynaptic and postsynaptic structures of the NMJ were enlarged in the BSG+GNP treated VML-injured muscles. This adaptation likely enables the NMJ to store increased amounts of ACh in synaptic vesicles and upregulate the expression of AChR receptors on the myofibers. These changes may help delay the onset of neuromuscular transmission failure, thereby mitigating the associated muscular fatigue. These findings suggest a protective effect and a positive overall impact of the BSG+GNP on both muscle and nerve repair. However, the extent of postsynaptic terminal fragmentation was increased despite BSG treatment, suggesting a lack of AChR organization, which will inevitably limit neuromuscular transmission and prevent functional recovery.

However, in the BSG+GNP+GDNF group, while the presynaptic terminals showed a rightward shift, indicating an increase in size, the postsynaptic terminal size was similar to that of the controls. Another important finding was the reduction in postsynaptic fragmentation with this treatment. The BSG+GNP+GDNF treatment strategy also resulted in increased co-localization between pre- and postsynaptic structures, as well as increased peak isometric torque production. Taken together, these findings suggest that the postsynaptic structures or AChRs were better organized and positioned to form a stable, accurate contact with the presynaptic terminal. A larger presynaptic terminal would suggest ample ACh release for effective signal transmission and torque production. Therefore, the improvement in NMJ quantity and function, driven by GDNF, provides the underlying mechanism for increased torque production in this group.

We speculate that an ECM-based BSG scaffold provides structural support and guidance cues for regenerating axons and myofibers while simultaneously modulating the inflammatory response, creating a conducive microenvironment for both muscle and nerve survival and growth [9–12, 27, 28]. However, this treatment alone was insufficient to enhance functional NMJ quantity. In other studies, improvements in functional NMJ quantity were not observed when BSG scaffolds were combined with eccentric exercise [27] or placental stem cells [26]. Alcazar *et al.* implanted aligned nanofibrillar collagen scaffolds loaded with insulin-like growth factor (IGF-1) into a mouse VML model. However, the NMJ quantity was increased only when scaffolds were combined with exercise [39]. In another study by Quarta *et al.*, decellularized bioconstructs containing muscle stem cells increased NMJ number only when combined with exercise. These observations suggest that incorporating potent neurotrophic or synaptogenic factors within scaffolds is necessary to enhance reinnervation and functional recovery.

Other groups have utilized biomaterials loaded with various biomolecules to both study and improve neuromuscular regeneration following VML. For instance, Mihaly *et al.* [40], and Scott *et al.* [41], used agrin-tethered fibrin scaffolds to enhance AChR clustering on skeletal muscle. Ziemkiewicz *et al.*, showed increased NMJ quantity and functional recovery in fibrin hydrogels supplemented with laminin-111 [42]. The success of these strategies further suggests that localized delivery of neurogenic signaling molecules is necessary for myofiber reinnervation. A key limitation of all the studies mentioned above is reliance on transverse 10–15 µm sections for quantitative analysis of NMJs. This method often yields inaccurate estimates of NMJ quantity and morphology. However, the preferred method of using longitudinal sections (>50 µm thick) for accurate NMJ analysis [43] is precluded in these experimental designs because multiple other measurements and analyses must be performed on the same tissue samples to determine the extent of tissue repair and regeneration.

The current study has several limitations. Although *in vivo* release kinetics were not measured, the enhanced recovery outcomes observed at 6 weeks post-injury in the BSG+GNP+GDNF group suggest prolonged GDNF availability. This sustained release is mechanistically supported by the slow degradation of the BSG, which was histologically identifiable even at 6 weeks. *In vitro* data corroborates this, showing an initial burst release of loosely bound GDNF within the first 3 days, followed by a slower, sustained release of electrostatically bound, deeply embedded GDNF within the BSG pores. Additionally, a single time-point (i.e., 6 weeks post-injury) was used for analysis. This prevents the assessment of the kinetic healing process. Furthermore, the need for thick longitudinal tissue sections for high-fidelity NMJ analysis, while beneficial for quantifying total NMJ count, precludes detailed analysis of muscle regeneration, specifically the quantification of myofiber CSA and fiber type distribution. Future studies should examine NMJ numbers at multiple time points and include transverse tissue sections to provide a more detailed analysis, thereby establishing a comprehensive, kinetic understanding of both muscle and nerve regeneration.

## Author Contributions

KG, PJ, MDW, and CRK designed the study. JT, CT, AR, KDS, RS, MEG, EGE, and KG performed experiments, collected data, performed statistical analysis, and prepared figures. JT and KG drafted the manuscript. All authors read and approved the final version for publication.

## Acknowledgements

The authors also thank Caroline Murphy at the SLU histological core for assistance and training with confocal microscopy.

---

**Supp. Fig. 1.**
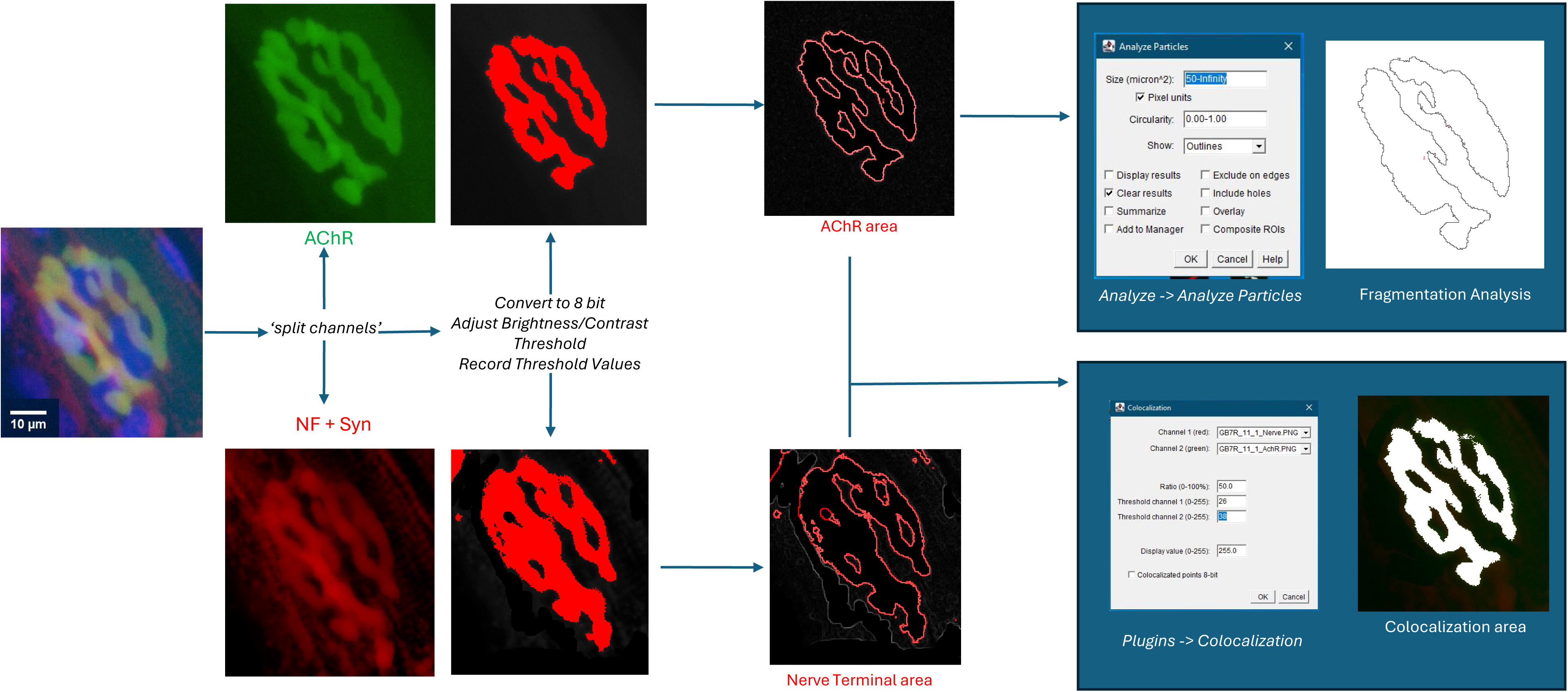
The quantitative analysis workflow utilized by blinded investigators is presented. A confocal image with merged channels is first split into individual channels, i.e., AChR and NF+Syn. The individual channels are then converted into 8-bit greyscale images, followed by adjustments to brightness and contrast. The channels are then thresholded, and the values are recorded. The threshold images are then used to measure the 2D-planar area of the individual channels, providing pre- and post-synaptic areas. The recorded threshold values were used to run the ’colocalization’ plugin on ImageJ to obtain the area of colocalization between pre- and post-synaptic structures. Finally, the analyze particles tool was used to obtain the number of fragments for the post-synaptic terminal.

**Supp. Fig. 2.**
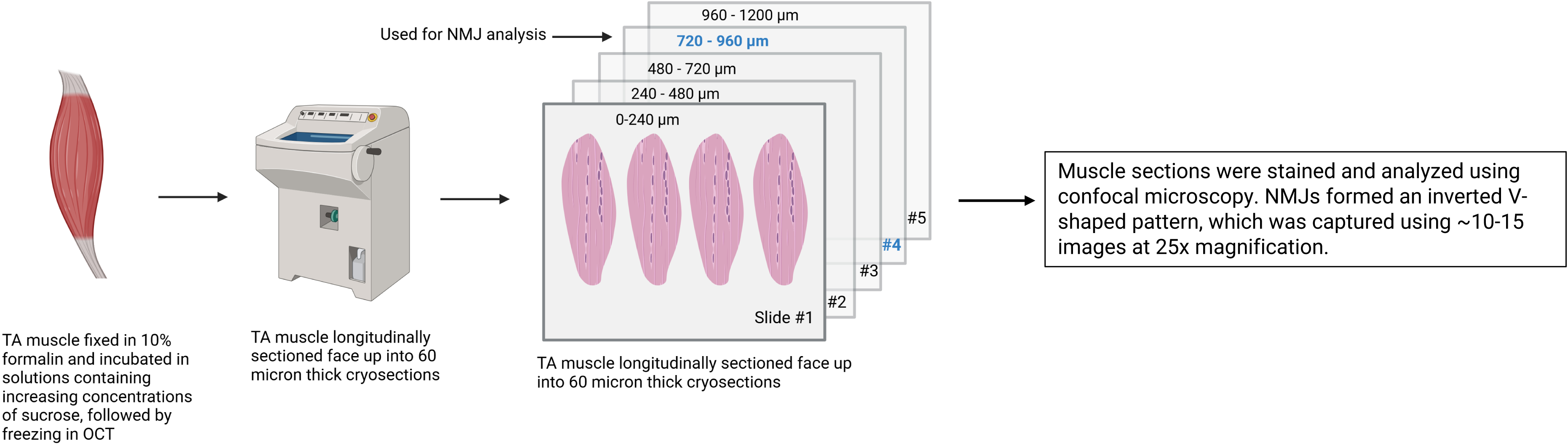
Post-harvest, the TA muscle was fixed in 10% formalin for an hour, followed by 30-minute soaks in increasing concentrations of sucrose solution (5%, 10% and 20%). The muscle was left in 20% sucrose overnight. The next day, the muscles were flash frozen in liquid nitrogen using OCT compound. The muscles were mounted face up on chucks and sectioned into 60 micron-thick sections. Muscle sections (four per slide) were mounted onto Superfrost^TM^ Plus Gold slides. Slides were numbered. Slides #4 and #5 were used for histological analysis.

